# Minuteman – A versatile cloud computational platform for collaborative research

**DOI:** 10.1101/2023.01.21.514916

**Authors:** Xinkai Li, Joydeep Charkaborty, Michael Jameson, Hobert Moore, Alexis Laux-Biehlmann, Sikander Hayat, Dhawal Jain

## Abstract

Secure platforms for bio-computation are critical to foster increasingly complex and data-intensive collaborations involving biomedical data. Here, we present Minuteman – an open-source cloud computing platform that can be securely used across organizations. Minuteman can be used for hosting data sources and running computational pipelines in an organized way. The platform consists of three fundamental features, 1) data operations including collaborative data processing and analytics, 2) customizable user access management for secure dissemination of the data, and 3) interactive exploration of the data through 3^rd^ party (e.g. shiny, dashboard etc.) applications that can be scaled using docker containers. Strict data access rules and user-specific roles are applied across the whole platform to maintain data security. Minuteman is ideal for scenarios where data security, and privileged access are critical, such as industry-academia collaborations, and multi-institution consortiums. Using single-cell transcriptomics preprocessing, analyses and visualization pipelines across labs, we showcase the utility of the Minuteman platform for biomedical data analyses. Minuteman code is available at (https://github.com/hayatlab/minuteman)

## 1. Introduction

An increasing number of industry-academia collaborations, multi-lab consortia and collaborations are needed to tackle some of the most complex challenges in the biomedical domain. For example, in the single-cell field, multiple labs[1], consortia such as the Human Cell Atlas (HCA)[2] and industry-academia collaborations[3] need extensive computational pipelines to process and analyze their biomedical data securely, and selectively share it across the collaborating organizations in real-time. This scenario needs a reliable computational platform, where participants can securely share their data and pipelines in a privacy-preserved manner. Several commercial solutions such as DNAnexus[4], SevenBridges[5], and Domino are available. However, they are expensive to run and maintain for a typical academic lab. Furthermore, due to proprietary nature, these platforms are not readily customizable by the end users. Here, we present Minuteman, a cloud-based computational platform-for collaborative data analyses. Minuteman can be used to perform various data operations using available as well as custom software and pipelines. Moreover, the same platform can be further used to disseminate the summarized data using any third-party data visualization solutions such as shiny, dashboard and django. The platform is built around cloud services provided by Amazon Web Services (AWS) where data can be securely shared in a privacy-preserved manner across collaborating organizations. Minuteman uses cloud-based parallel computing for resource intensive data operations, R/Python or similar environment for focused data analytics and shiny/dashboard and django framework for data visualization. The userinterface of Minuteman is developed in Django and can be accessed through weblink upon installation. The capabilities of Minuteman platform allow users to manage data resources and codes in an organized manner hence, supporting the “Findable, Accessible, Interoperable and Reusable” (FAIR) principles of data operations. Here, we describe Minuteman and its utility in analyzing single-cell transcriptomics data. We believe that Minuteman will be a useful platform for computational analyses and will foster collaborations.

## 2. Results

### 2.1 Architecture of Minuteman

Minuteman is built using Amazon Web Services (AWS). The Elastic Kubernetes Service (EKS), AWS ParallelCluster, Django, Jupyterhub and Rstudio community edition are its foundational building blocks. However, the servers within this framework can be further customized by the users according to their requirements. Minuteman runs inside the virtual private cloud (VPC) of AWS with two distinct subnets - private and public (Figure 1). The private subnet refers to the organization hosting the infrastructure, while collaborating labs are referred to as public. Traffic to private subnets is blocked from the public network and regulated from public subnets to allow restricted and customizable access. Resources such as high-performance clusters (HPC) for parallel computing, various relational databases, data analytic environments such as R/Python, data visualization containers are hosted inside the private subnet. Internal users, from the hosting organization, can reach the private subnet directly via organizational VPN and through user authentication system. Public subnets on Minuteman can be reached from public networks via the secure gateways and user authentication system. These gateways can be configured to whitelist the incoming traffic to ensure maximum data security. All incoming traffic to Minuteman is gated through load-balancers for optimal performance. The EKS is used to host website infrastructure that further directs users to various containerized applications or services for data analytics or data processing (Figure 1). For instance, Minuteman uses AWS products such as Parallel Cluster with slurm engine for high performance computing, Amazon Relational Databases with SQL servers for holding various data, Glue, Athena and S3 for ETL pipelines. We use Gitlab as code repository while gitlab runner and custom bash scripts for continuous integration continuous delivery (CICD). Data is stored on S3 buckets that can be made available to the private subnets. Overall, the architecture of Minuteman is modular and new features such as third-party data analytics/visualization software or Amazon products can be easily integrated into the private subnets.

**Figure 1.**
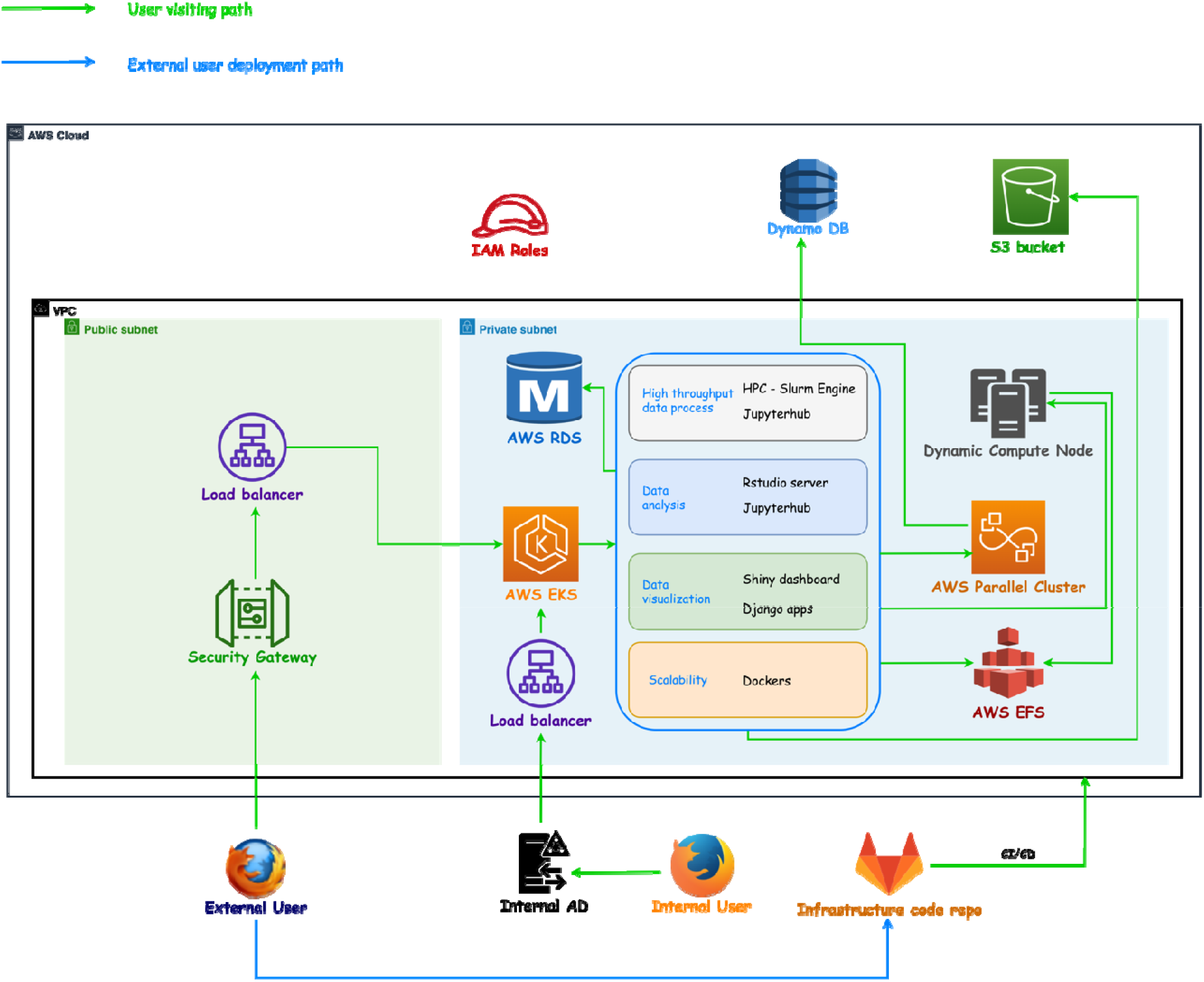
Overview of the Minuteman platform. The platform is organized into private (internal or hosting) and public (external or collaborating) subnets. Data processing, visualization and interface infrastructures are located inside a private subnet. Users from collaborating organizations are routed through secure gateways to the private subnet. Various Amazon services are used for optimal performance of the infrastructure. The private subnets are highly customizable where data analytic environment, tools or other Amazon services can be integrated.

### 2.2 Computation and job management

Minuteman provides computational capability for running datasets of various dimensions. It uses dynamic compute nodes based on AWS ParallelCluster for running high performance computing applications. The slurm engine is used to submit parallel jobs. For each job, the scheduler spins an EC2 instance, monitors the job progress and shuts it down when the job is completed. For relatively light containerized or data visualizations applications, Minuteman uses horizontal pod autoscaling. When the CPU usage exceeds 50% the system generates replica pods, until the usage is below 50%. Minuteman uses pods with stateless nginx gateways for autoscaling (Figure 1).

### 2.3 Data security and privacy preserved access

Minuteman organizes data storage in three different ways - S3 buckets, RDS/SQL databases and local storage. Overall access to the infrastructure and underlying data is typically managed at the user level at various levels. First, as the infrastructure is built on a private subnet, user access is established under the VPN or VPC of the hosting organization. Second, users can access data on Minuteman only through their user profiles -limiting access in user home or shared spaces. And third, data stored in RDS/local are accessible in a role-based manner. AWS secret manager is employed for password protecting the databases. For users of various applications, user access is managed separately. Users belonging to the hosting organization (internal users) are validated through the host userbase. And the users from collaborating organizations (external users) are validated using a Django framework. Various user groups (currently superuser and user) are defined and access is managed using the same Django framework (Figure 2).

**Figure 2:**
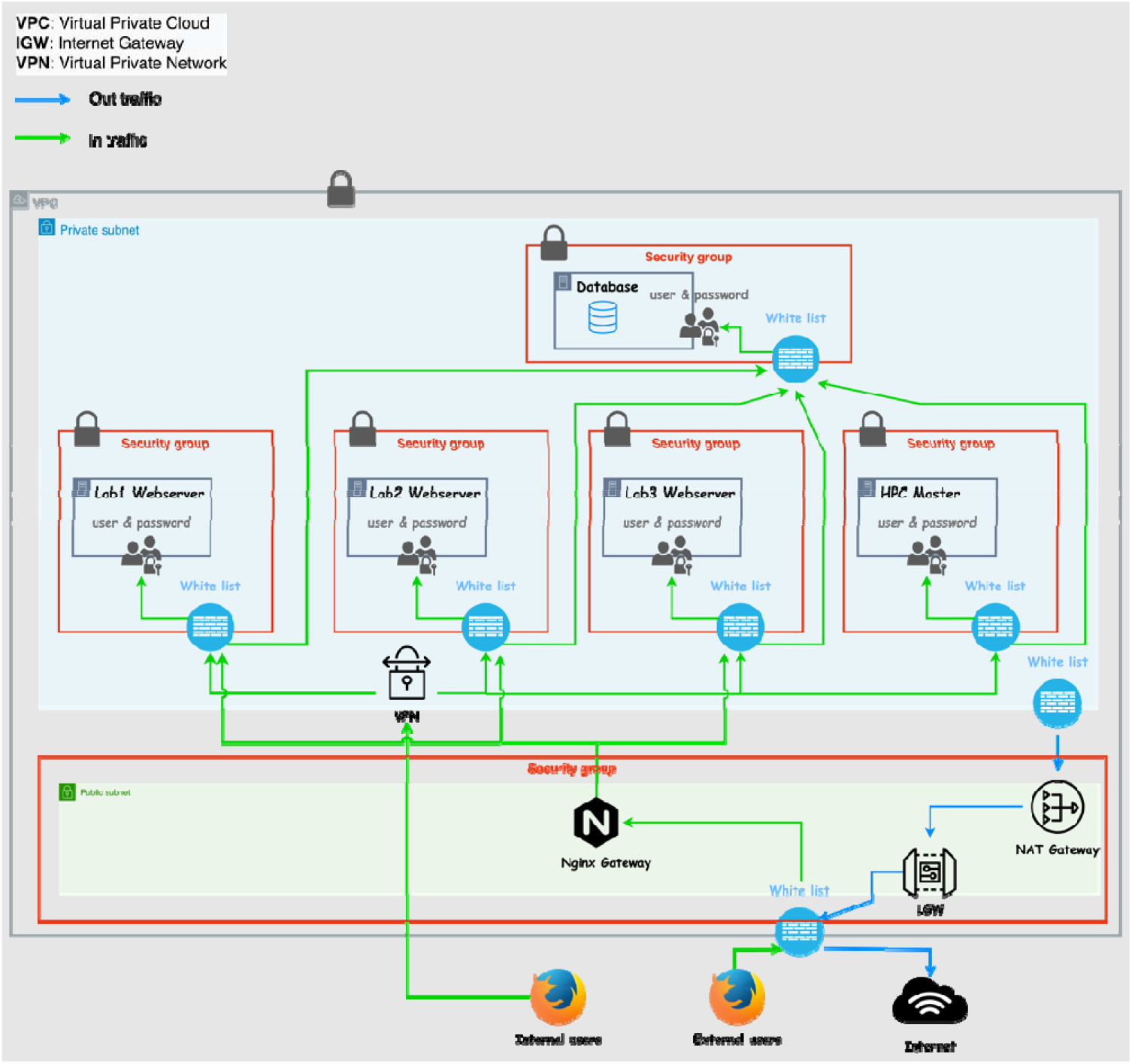
Minuteman security and firewall infrastructure. Several research collaborations can be managed in Minuteman as separate private networks that can be completely isolated from other private networks of the hosting subnet. Here, user access and roles are maintained separately. Both internal and external users can be granted access to the VPN securely.

### 2.4 Devops and collaboration

Oftentimes, collaborations result in the development of new data analytic or visualization applications. These applications can be shared by containerization across the collaboration on Minuteman. For instance, the application repo on Gitlab can be used by the development teams to jointly develop applications. Once new tags are created, a CICD is triggered to build a new docker image with an updated tag. These new images are then deployed on the deployment repo and hence on Minuteman infrastructure under the appropriate private subnet. This design can easily realize collaboration among teams across organizations. The gitlab repo can also be used to upload separately developed applications in the form of docker images. (Figure S1).

## 3. Case study: Utility of Minuteman for single-cell transcriptomics data analyses

We extensively used the Minuteman platform for running single cell data processing and visualizations (Figure 3; Figure S2). The data processing pipeline uses *mkfastq* and *count* functions from cellranger[7] to generate a count matrix from raw bcl data. The pipeline was containerized and run on Minuteman using parallel computing. The-UMI count matrices generated with the processing part were then aggregated and passed as input to our custom QC pipeline that filters out cells and genes based on factors such as reads mapped to mitochondrial genes, number of cells expressing a gene. The cleaned matrix was then passed to the pipeline for batch effect correction. Here, we have currently implemented Harmony[8], Seurat[9] and scVI[10] for batch effect correction and calculation of a low-dimensional representation of the data using highly variable genes. This low-dimensional representation was then used for clustering, marker gene detection, and subsequent downstream analyses such as compositional changes, differentially expressed genes and dysregulated pathways discovery (Figure S2). Lastly, we used CellXgene[6] and SciView[11] applications for interactive visualization of the single cell sequencing datasets on Minuteman.

**Figure 3:**
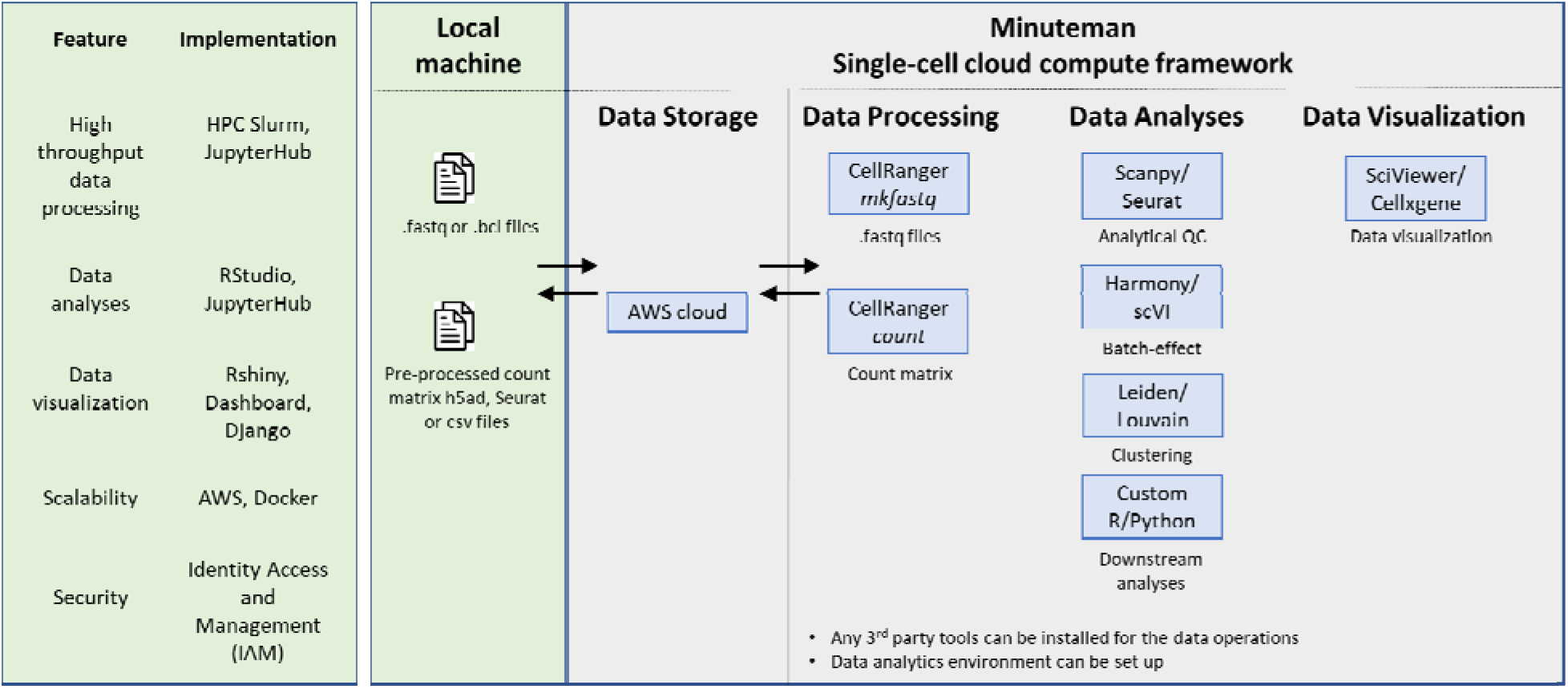
Overview of Minuteman for single-cell transcriptomics analyses. Raw data files such as *fastq* and *bcl* files, or preprocessed count matrix objects can be uploaded to the Minuteman cloud framework. Computational pipelines available in Minuteman allow for data preprocessing (using cellranger), data analyses (Scanpy and Seurat), and downstream analyses using custom R and Python scripts in a scalable manner using AWS cloud compute framework. Interactive data visualization applications such as Sciviewer and Cellxgene[6] can also be deployed in Minuteman. The modular architecture of Minuteman allows users to develop their own custom applications for data exploration.

## 4. Discussion

We present here – Minuteman, a computational platform to collaboratively work on biomedical datasets of high value-. Minuteman stems from our real-world experience of working with high value single-cell transcriptomics and other datasets across different labs in both industry and academia. The platform offers possibility for computational scientists to collaboratively analyze the data and disseminate the synthesized information to the target audience of bench scientists by either building custom visualizations or using preexisting data visualization tools. All these operations can be securely performed on Minuteman. The platform can be used to collaborate with multiple labs where data sharing is highly secure and achieved in a role-based manner. The Minuteman infrastructure is modular such that any new software, data analytical environment or collaborative workspace can be added to it. With ease of sequencing and multidimensional nature of scientific discourse, the need for collaborative research is growing rapidly. In this data-driven collaborative landscape, we believe, Minuteman will be a useful open-source platform that will not only foster collaborations between different labs but also bridge the gap between computational and biology researchers.

## Supplementary Data

### Guide to install and deploy Minuteman

A step-by-step guide to install and deploy Minuteman is provided with the Minuteman codebase available at: https://github.com/dhawalsjain/SciViewer

a. Build basic AWS infrastructure
  i. Build VPC
  ii. Build keypairs, IAM roles, security groups
  iii. Deploy gitlab-runner and connecting it to gitlab repository
  iv. Deploy EKS and kube-dashboard, metrics-server
  v. Deploy MySQL on RDS
  vi. Setup route 53
  vii. Apply domain and SSL certificate
b. Build HPC
  i. Build EFS
  ii. Deploy main AWS Parallel cluster
c. Deploy main website
  i. Deploy main website through CICD to EKS
d. External user setup
  i. Deploy AWS ELB on a public subnet.
  ii. Setup related security groups

### Supplementary Tables

**Table S1:**
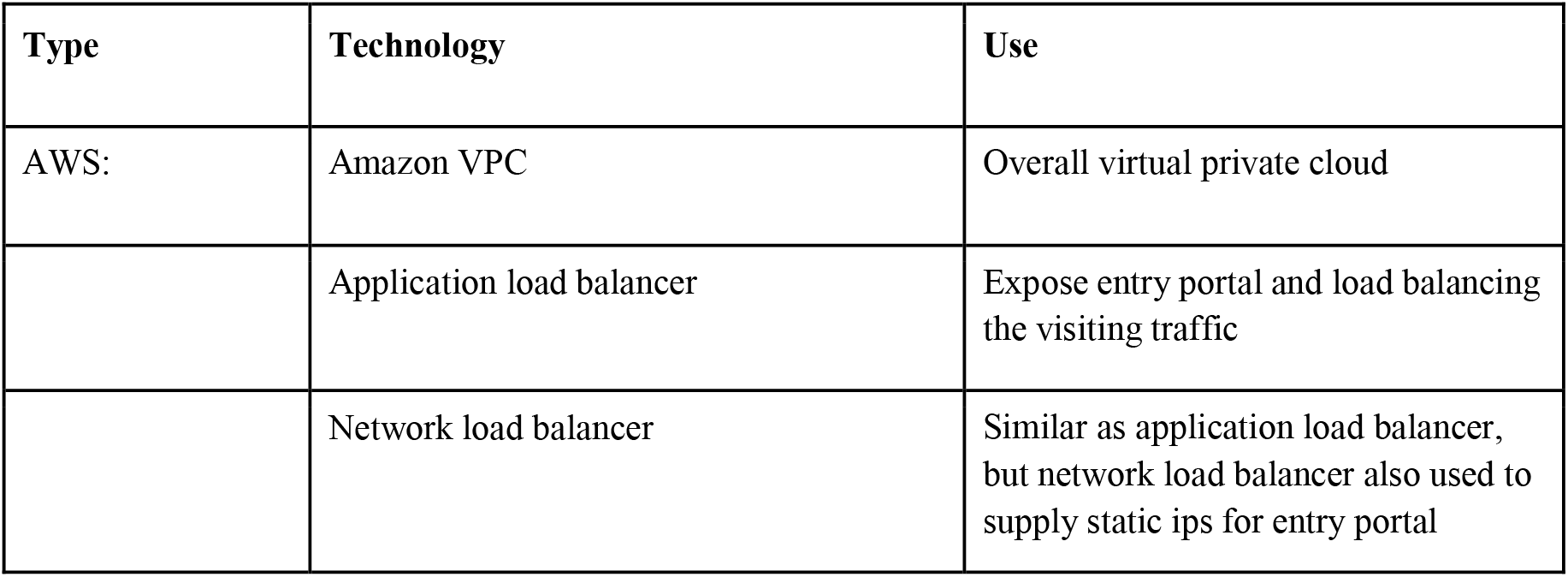
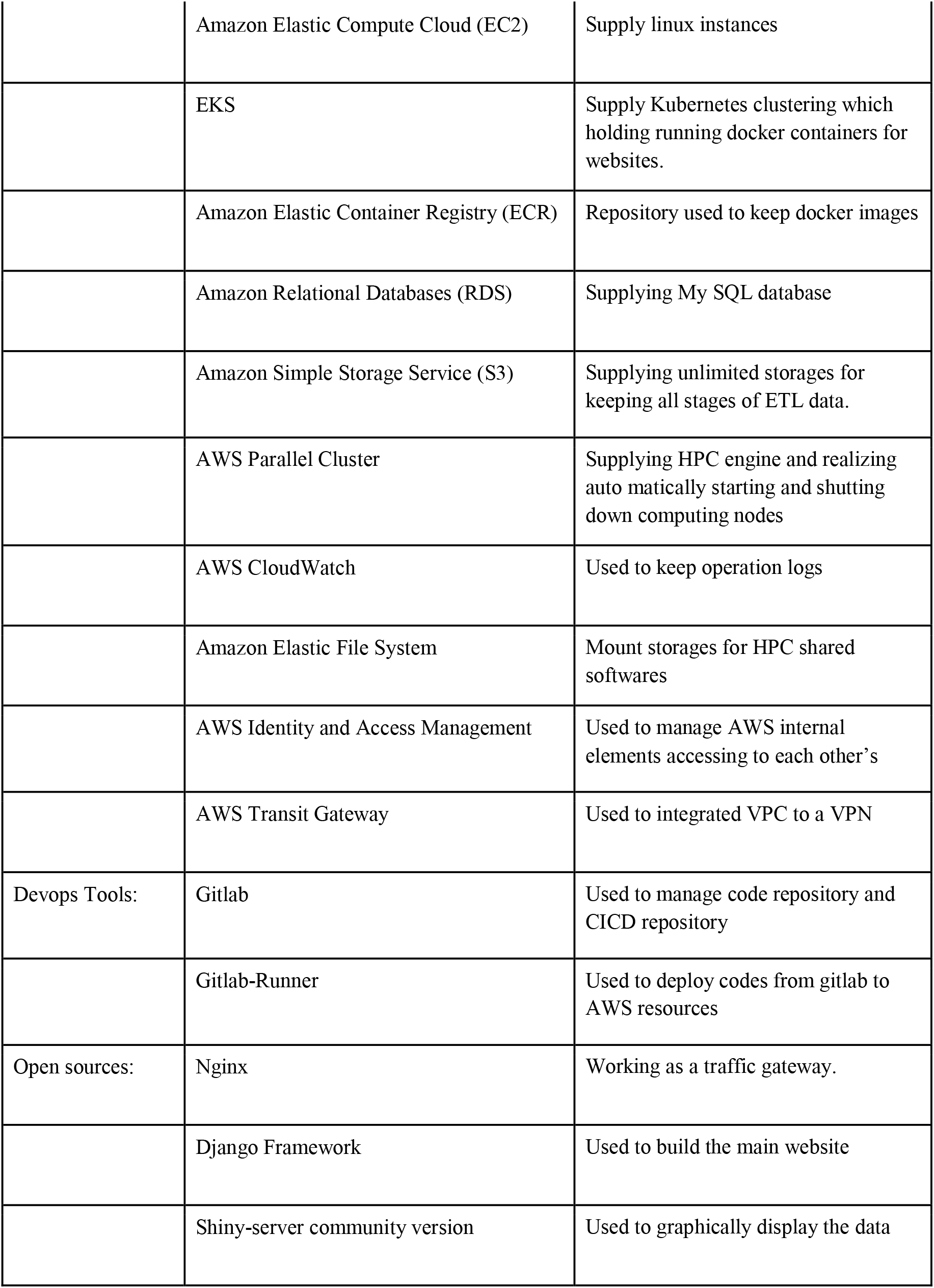
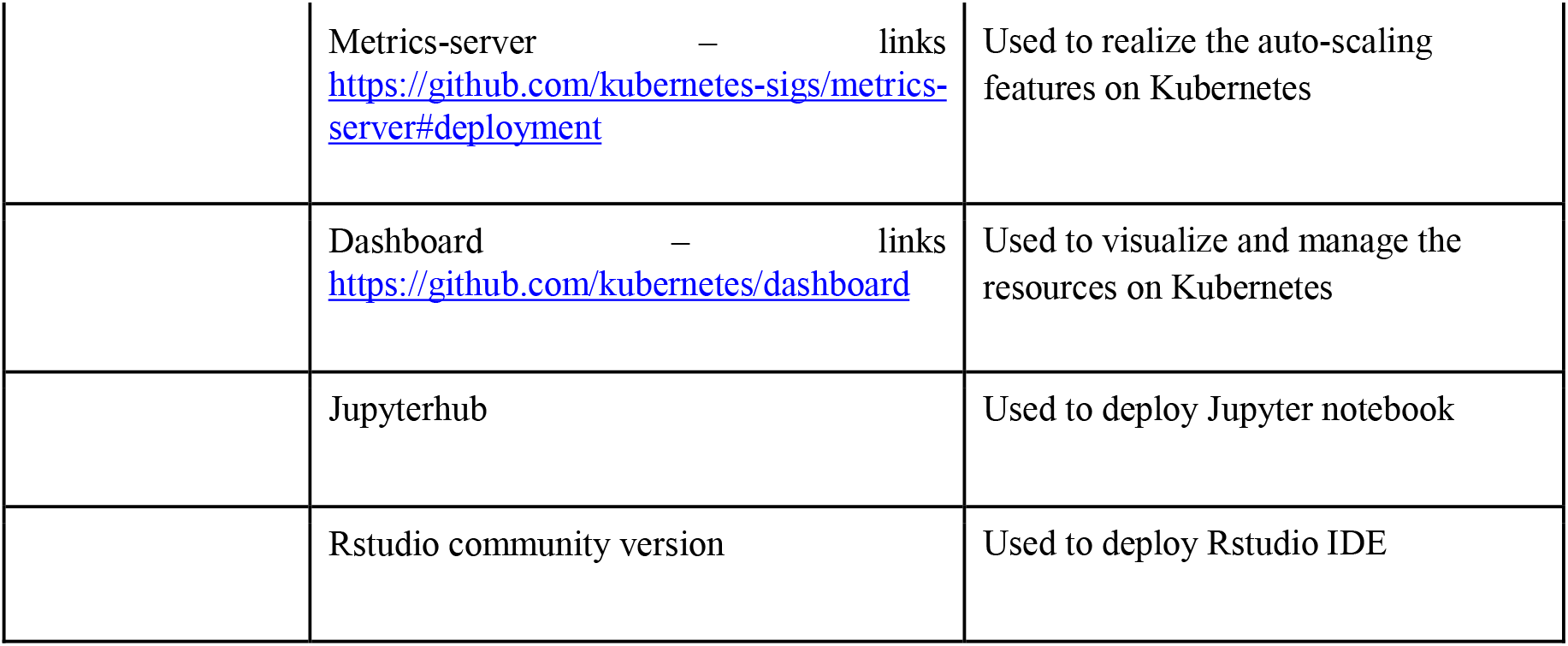
List of all technologies that were used to develop and implement Minuteman.

**Figure S1:**
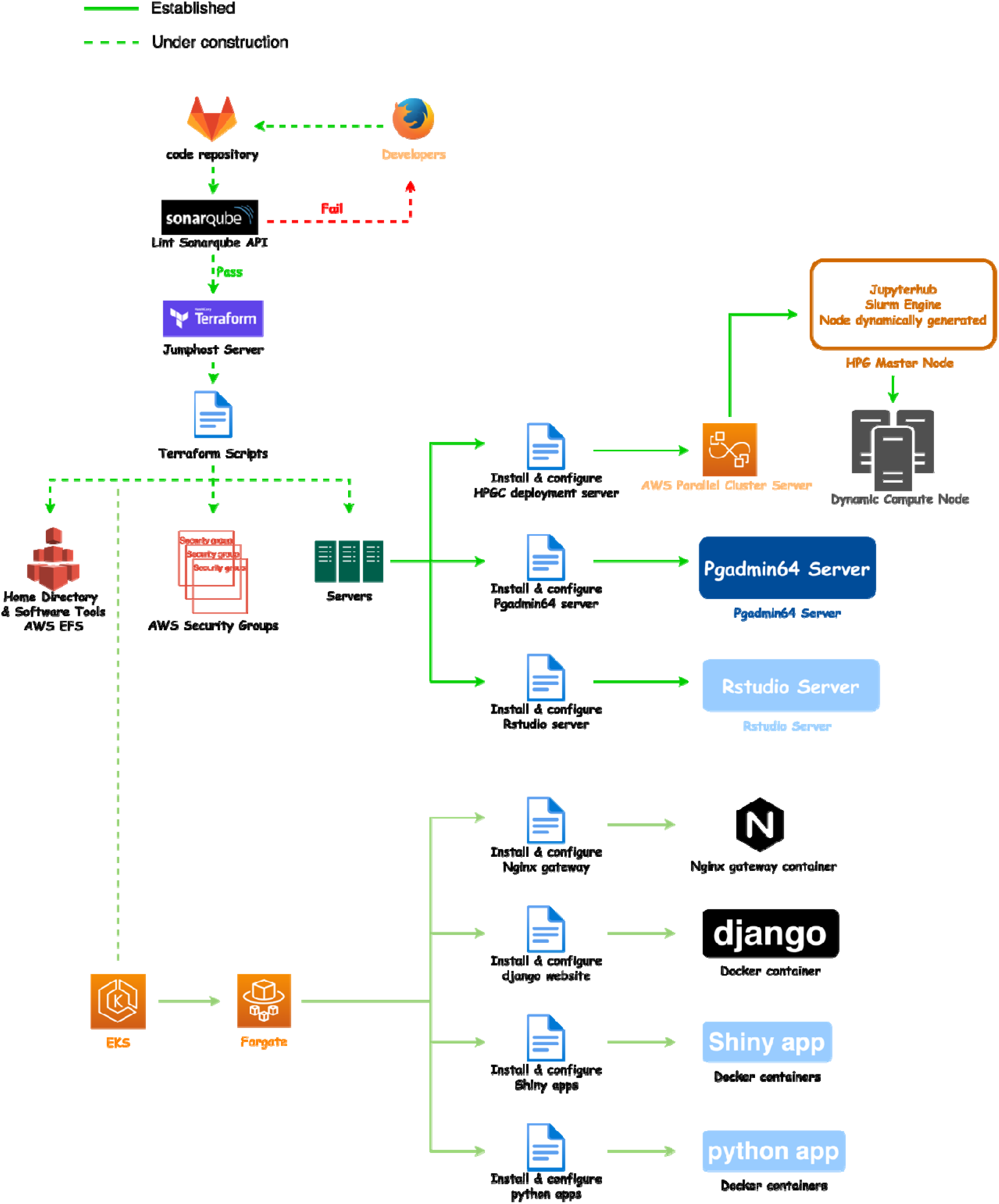
Overview of workflows that can be deployed using Minuteman. Detailed overview of established workflows that have been implemented in Minuteman. New features, such as, tools for code maintenance and security that can be readily implemented within the Minuteman framework.

**Figure S2:**
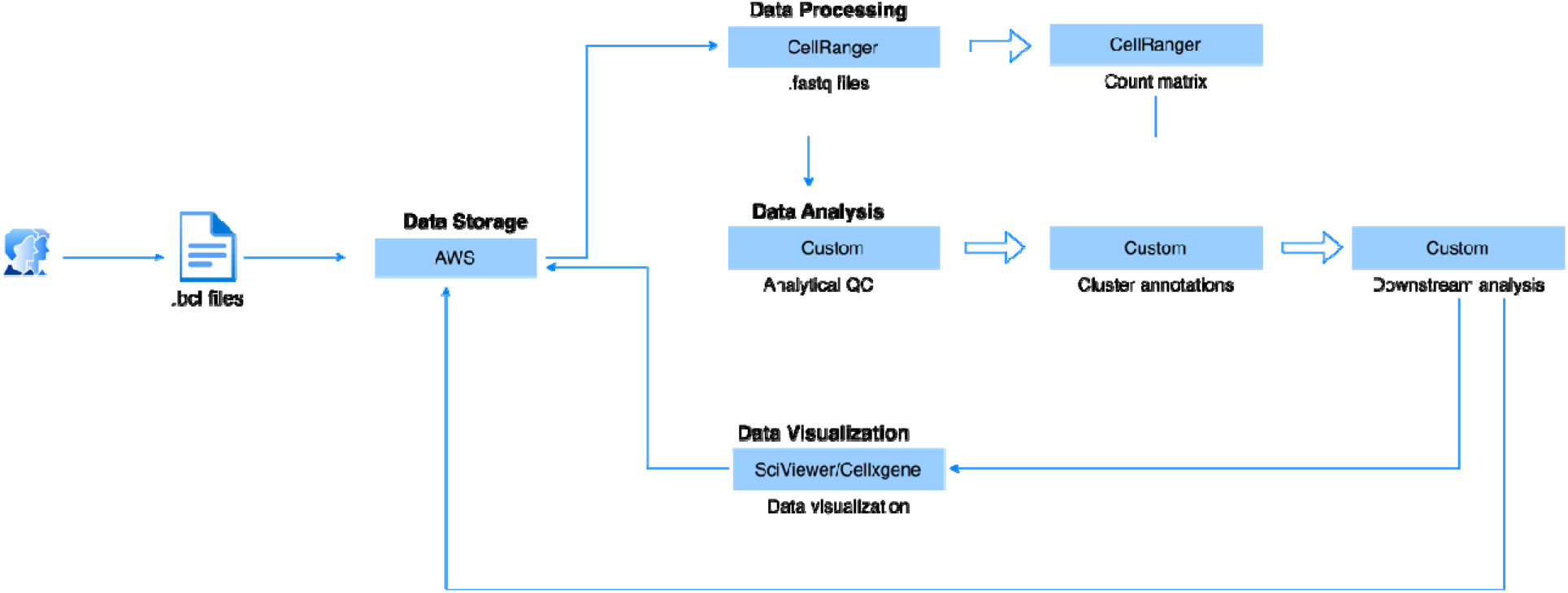
Implementation of a single-cell transcriptomics pipeline for data processing, analyses and visualization using the Minuteman framework. Here, data objects can be securely shared, and pipelines can be scaled based on the computational needs.

